# Immediate reuse of patch-clamp pipettes with sonification

**DOI:** 10.1101/2022.11.11.515952

**Authors:** Kevin Jehasse, Sophie Wetz, Peter Neumann-Raizel, Björn M. Kampa

## Abstract

The patch-clamp technique has revolutionized neurophysiology by allowing to study single neuronal excitability, synaptic connectivity, morphology, and the transcriptomic profile. However, the throughput in recordings is limited because of the manual replacement of patch-pipettes after each attempt which are often also unsuccessful. This has been overcome by automated cleaning the tips in detergent solutions, allowing to reuse the pipette for further recordings. Here, we developed a novel method of automated cleaning by sonicating the tips within the bath solution wherein the cells are placed, reducing the risk of contaminating the bath solution or internal solution of the recording pipette by any detergent and avoiding the necessity of a separate chamber for cleaning. We showed that the patch-pipettes can be used consecutively at least 10 times and that the cleaning process does not negatively impact neither the brain slices nor other patched neurons. This method, combined with automated patch-clamp, highly improves the throughput for single and especially multiple recordings.

## Introduction

Patch-clamp recording is a widely-used and powerful technique to study single-cell electrophysiology, especially in neuroscience. It led to the characterization of several aspects of neuronal physiology, *in vitro* and *in vivo*, such as ion channel activity underlying action potential^1,2^, intrinsic excitability^3,4^, synaptic integration^5,6^ plasticity^7,8^ and network activity^9,10^. Patch-clamp also allows recordings in the different compartments of neurons^11–13^ including very distant dendritic branches or axonal sections^14^. In the *whole-cell* configuration, it is possible to dialyze cells with fluorescent dyes for morphological reconstructions and to collect the cytoplasm to analyze single-cell transcriptomic profiles^15,16^. Although the patch-clamp technique is well suited to characterize the heterogeneity of neurons in the brain, it is highly laborious and time-consuming with variable success rates resulting in a low-throughput. To overcome this issue, engineering advancements managed to automate patch-clamp *in vitro* and *in vivo*^17–19^. When automated, the software is able to track the patch pipette and individual neuron positions in order to approach and record them. While it reduces the human interaction for the recording, it is still required to manually change the patch pipette after each attempt.

To obtain a successful recording, one crucial parameter is to have a perfectly clean patch-clamp pipette tip filled with a filtered internal solution^20^. In this condition, there is a high chance to form a seal of high-resistance (≥ 1 GΩ, which is called “gigaseal”) with the membrane of the neuron. After each successful or failed attempt, the used pipette has to be manually changed. Therefore, the human factor can be a big issue when it comes to multiple recordings because of vibrations from the exchange that can affect the seal of other pipettes attached to neurons in the context of neuronal network characterization. It can also be an issue for *in vivo* patch-clamp studies as the manual pipette replacement could disrupt the animal and the following recordings. To prevent that, an automated method for cleaning patch pipettes has recently been developed^21^. This approach consists of immersing the tip of used patch-clamp pipettes in a separate chamber containing the detergent Alconox while applying cycles of positive and negative pressure to the pipette tip. Coupled with automated patch-clamp^22^, the throughput of patch-clamp recording is significantly increased. However, the necessity of a separate bath cleaning chamber and the risk of contaminating the recording bath solution nor pipette internal solution with detergent reducing the usability of this cleaning method.

Since patch-clamp pipettes are made of borosilicate glass and can also be cleaned by sonification, we developed a novel cleaning system based on that method preventing the use of detergent. The sonification is performed by a piezo-element mounted on the recording headstage and connected to a pressure control system (Fig.1a). The ultrasonic cleaning is performed in artificial cerebrospinal fluid (aCSF) and allows to reuse the same patch-clamp pipette at least 10 times without affecting the recording quality. We also showed that the cleaning procedure within the same bath does not affect the stability of seal of other patch-clamp pipettes with their respective neurons. Therefore, ultrasonic cleaning is a powerful improvement offering significant advantages in particular for multiple simultaneous patch-clamp recordings.

**Figure 1:**
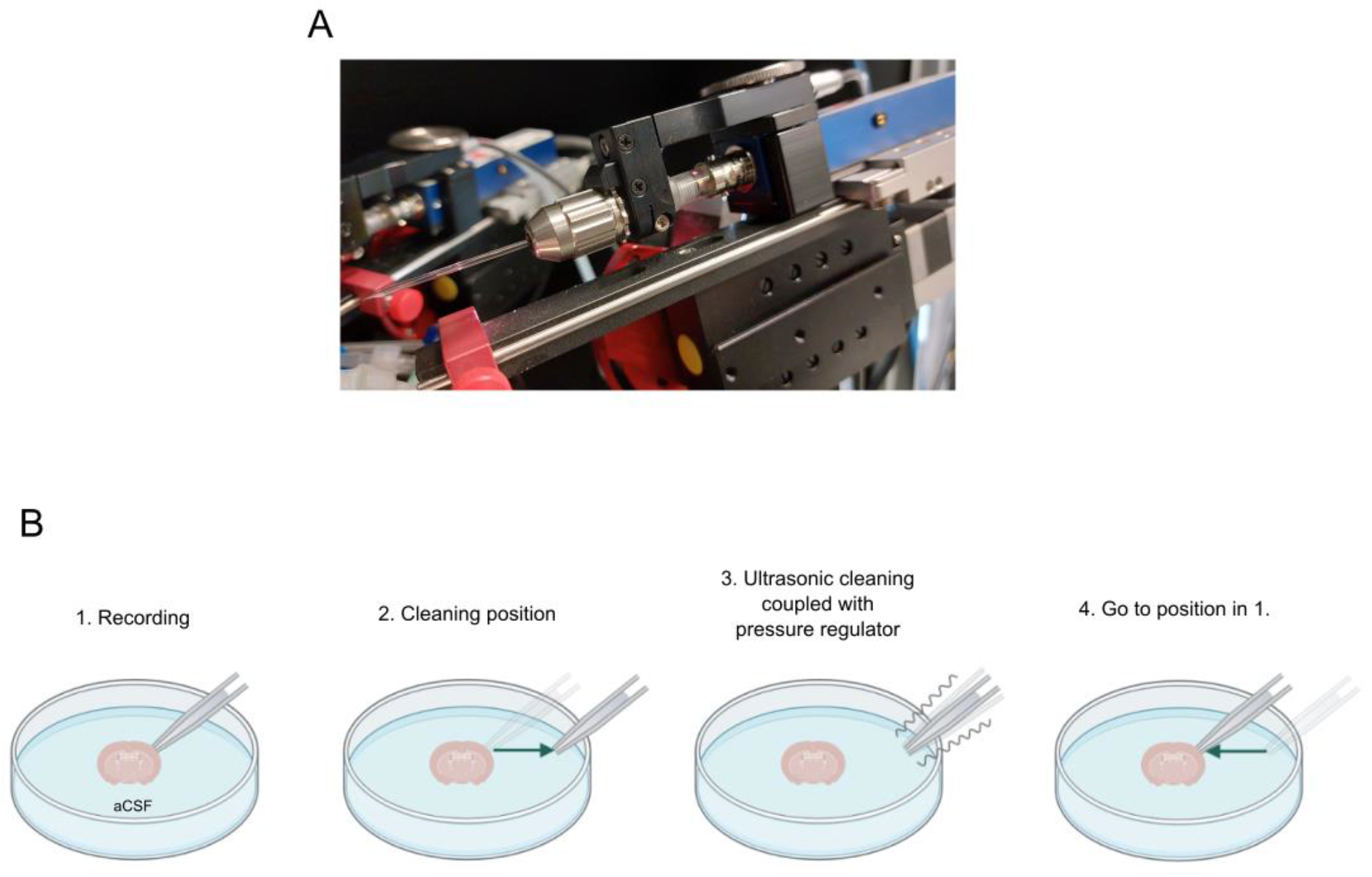
Ultrasonic cleaning is performed by a piezo coupled to a pressure regulator. **(A)** Picture of the headstage unit. This unit is connected to a pressure regulator to allow the cleaning. **(B)** When the recording is finished (1), the cleaning sequence can be launched. The manipulator put the tip to a pre-defined cleaning position (2) where the sonification coupled with cycle of high positive and negative pressure can occur (3). Once the procedure is finished, the manipulator put the tip back to its initial parking position (4) before the cleaning was launched.

## Results

### Cleaning efficiency

Before approaching the patch-clamp pipette to the brain slice, a “cleaning position” (> 5 mm from the slice) was defined to safely clean the pipette without affecting the sample. This is determined by the micromanipulator control panel. When “cleaning” is selected, the micromanipulator automatically goes to the saved XYZ coordinates (Fig. 1B). The piezo mounted on the headstage performs the sonification of the patch-clamp pipettes by making it oscillate at 40 kHz. To help removing the membrane residue inside the pipette, the sonification is coupled with a cycle of high positive and negative pressure similar to the protocol used for cleaning in detergent^21^. After the cleaning procedure the patch-clamp pipette can be placed into a “parking position” which correspond to a determined XYZ distance to its original place. This prevents any damage of the slice by the automatic replacement of the pipette and to freely move to find another neuron to record.

For each attempt, the pipette resistance (R_pip_) was monitored once a *whole-cell* recording attempt was made, and compared before and after the ultrasonic cleaning (Fig. 2A) to assess the efficiency of the procedure. We used pipettes with R_pip_ between 3-5 MΩ and between 8-15 MΩ to test the efficiency of ultrasonic cleaning on patch-pipettes with regular tip size and sharp tip size regularly used, respectively for somatic recordings and for dendritic recordings^23^ or *in vivo* patch-clamp. As expected, R_pip_ was reduced after sonification (before: 13.97 ± 0.45 MΩ; after: 6.75 ± 0.04 MΩ, n = 88 attempts, p < 0.0001, Wilcoxon signed-rank test). Over ten cleaning cycles, R_pip_ remained constant (p = 0.6454 for fresh vs tenth cycle, n = 8, Wilcoxon signed-rank test; Fig. 2B). These data show that ultrasonic cleaning coupled with alternating high and low pressure can successfully remove remaining membrane residues and repeatedly over at least ten attempts.

**Figure 2:**
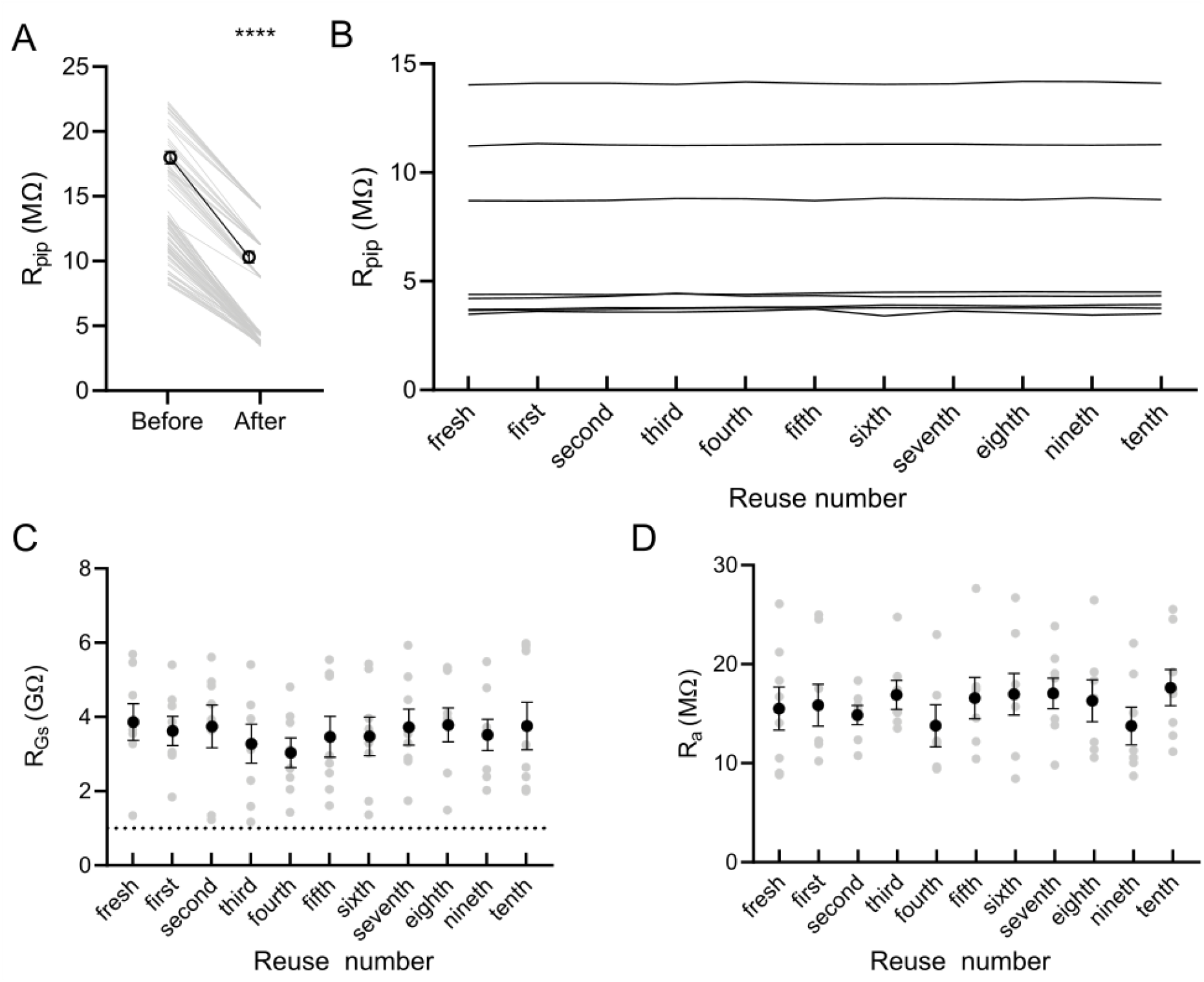
Ultrasonic cleaning allows reusing pipettes for multiple consecutive recordings. **(A)** and **(B)** indicate that ultrasonic cleaning can clean the tip as R_pip_ is recovered after each cycle. **(C)** shows that patch-clamp pipette can be successfully reused at least ten times and R_Gs_ is successfully reached. **(D)** show that R_a_ remains unaffected after each cleaning cycle, indicating that the tip is cleaned and the pipette can be reused for successful patch-clamp recordings.

Next, we evaluated the quality of recordings by monitoring the resistance of gigaseal (R_GS_) and access resistance (R_a_). R_GS_ was obtained before breaking in *whole-cell*, when the value was steady. We managed to reach gigaseal in 88 neurons and *whole-cell* configuration in 81 neurons with 8 pipettes (92.05% success rate) that were cleaned 10 times and we observed no effect of ultrasonic cleaning on R_GS_ (p = 0.988, Friedman test; Fig. 2C) and R_a_ (p = 0.8192, Kruskal-Wallis test; Fig. 2D). These results show that with the novel sonification method patch-clamp pipettes can be reused at least 10 times for successful *whole-cell* recordings.

### Effect of sonification on cell survival

We next tested whether ultrasonic cleaning could affect neuronal survival within the same chamber. In the context of simultaneous paired recordings, it is crucial that cleaning one pipette tip does not affect other neurons maintained in *whole-cell* by the patch-clamp pipettes. As mentioned above, the pipette is put at a safe distance (>5 mm) from the sample to prevent any harm or energy transfer to the other pipettes. In this condition, the resting membrane potential (RMP) of neurons being recorded simultaneously remained stable during the cleaning process (before: -65.0 ± 1.8 mV; after: -65.1 ± 1.9 mV, n = 5, p = 0.5625, Wilcoxon signed-rank test; Fig. 3A, B) and membrane resistance (R_m_) is not affected (before: 110.3 ± 9.3 MΩ; after: 109.7 ± 7.7 MΩ, n = 5, p > 0.99, Wilcoxon signed-rank test; Fig. 3A, C). In one example case, we successfully recorded the same neuron 3 times with the same patch-clamp pipette after being cleaned and filled it with biocytin for confirmation (Fig. 3D, E). Its intrinsic parameters remained stable for the all 3 attempts. And the post-hoc reconstructed morphology did not show any sign of neuronal damage (Fig. 3E). Altogether, these data show that ultrasonic patch-clamp pipette cleaning does not harm neurons situated within the same bath chamber.

**Figure 3:**
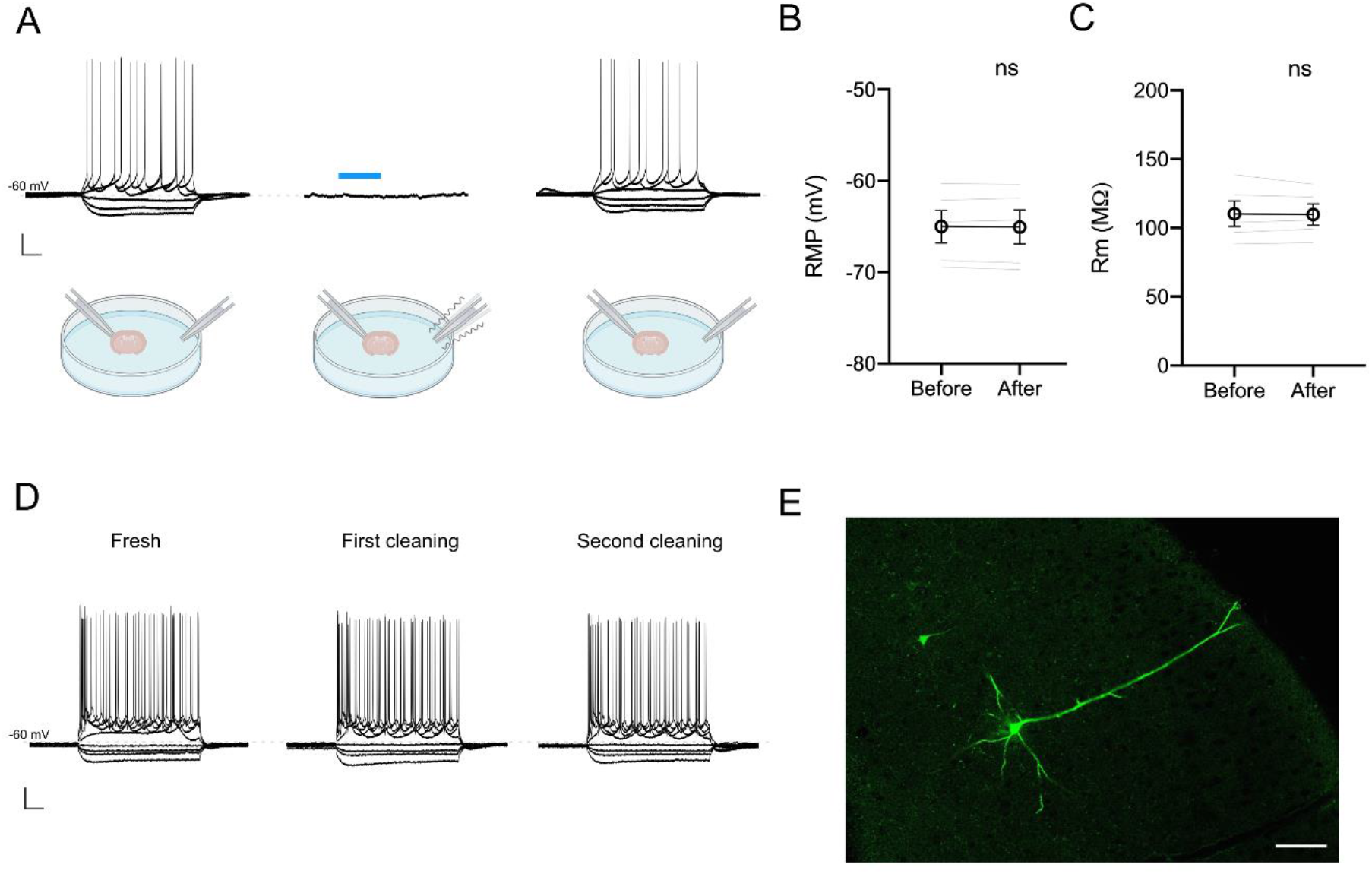
Ultrasonic cleaning is not harmful for neurons. **(A)** Example of a neuron being recorded before (left), during (middle) and after (right) ultrasonic cleaning of a patch-clamp pipette inside the same recording chamber. Scale bar is y = 20 mV (all panels) and x = 100 ms (left and right) and 20 s (middle). Blue bar corresponds to the moment when the non-recording tip is being cleaned. **(B)** and **(C)** show that the cleaning procedure does not interfere with simultaneous patch-clamp recordings as RMP and R_m_ of the recorded neurons remain unaffected. **(D)** Example a neuron being successfully recorded three consecutive times with the patch-clamp pipette after ultrasonic cleaning. Scale bar is y = 20 mV and x = 100 ms. **(E)** *Post-hoc* staining of neuron from **(D)**. Scale bar is x = 80 µm.

## Discussion

The patch-clamp technique is a very powerful method but it is laborious and the success rate is not high. In order to increase its throughput, automatization of the patch-clamp technique has recently been the focus of engineering. While there have been several successful developments to automatize the whole procedure^17,19,22^, it is crucial to overcome the changing of patch-clamp pipette. Others have shown that it is possible to clean the tips using either bleach^24^ or detergent combined with automated pressure control^21,25^. However, even if not detected, there might be some trace left of these chemicals inside the patch-clamp pipette that can be harmful for the cells. We developed an ultrasonic cleaning system coupled with automated pressure control, which allows the tips to be cleaning in a physiological solution and is less likely to harm the cells.

Cleaning patch-clamp remains to this day one of, if not, the key element to make patch-clamp a routine. In the context of drug discovery, automated patch-clamp experiments are essential to explore the pharmacology of ion channels. Although we did not investigate the effect of ultrasonic cleaning on ion channels, Kolb and colleagues showed that GABA receptors pharmacology is not affected with pipettes being cleaned in Alconox^21^. With these observations combined with ours, showing that ultrasonic cleaning does not affect neuronal excitability, it is highly possible that our method is not affecting the pharmacology of ion channels either.

Regarding the longevity of pipettes, we did not investigate how many times pipettes can be used before replacing it with a fresh one. Kolb and colleagues showed that it is not possible to reuse the same pipette indefinitely^21^. We suspect that our approach will allow reusing the same pipettes for a limited time, as internal solution contains chemicals that degrade when not refrigerated (e.g. ATP, GTP). Yet, for the time tested in our experiments, we could repeatedly record even from the same neurons for up to 10 times suggesting that the life time of the pipette is not a limiting factor.

By being able to clean the pipette tip in standard aCSF, our method offers the possibility to avoid using a second compartment. This is convenient, especially for *in vivo* patch-clamp, as it would allow cleaning the pipette above the exposed brain and reduce the time required to change it after each attempt. Coupled with recently automated *in vivo* patch-clamp recordings^17^, it will increase the throughput of these recordings in vivo and make the method also more accessible.

In conclusion, we have developed a detergent free cleaning method for patch-clamp pipettes based on sonification coupled with a pressure control system that can be implemented for automated patch-clamp systems.

## Methods

### Cleaning procedure

We used a pressure control system (LN-PCS, Luigs & Neumann, Germany) to deliver positive and negative pressure to the patch-pipette. The procedure is set and triggered with a SM10 Remote Control Touch (Luigs & Neumann, Germany). The cleaning protocol consisted of 6 steps alternating positive (+500 mBar for 3s) and negative pressures (−300 mBar for 3s) and the frequency of sonification is set at 40 kHz. Different pressures or frequencies can be used. A safe cleaning distance (> 5 mm from the brain slice) was defined before the beginning of the experiment to prevent any energy transfer from the pipette being cleaned to other pipettes attached to neurons in *whole-cell* configuration.

### Electrophysiology

All procedures were performed with the approval of local authority (LANUV NRW, Germany). Acute brain slices were obtained as previously described^26^: adult C57BL/6 mice of both genders were anesthetized with isofluorane (AbbVie, UK) and decapitated. 300 µm-thick coronal slices were cut with a vibratome (Leica VT1200s) in an ice-cold modified cutting solution containing (in mM): 125 NaCl, 2.5 KCl, 1.25 NaH_2_PO_4_, 25 NaHCO_3_, 25 Glucose, 6 MgCl_2_, 1 CaCl_2_, pH 7.4 (95% O_2_ / 5% CO_2_ and 310 mOsm/l). Slices were incubated for 30 min at 34°C in aCSF containing (in mM): 125 NaCl, 2.5 KCl, 1.25 NaH_2_PO_4_, 25 NaHCO_3_, 25 Glucose, 1 MgCl_2_, 2 CaCl_2_, pH 7.4 (95% O_2_ / 5% CO_2_ and ∼310 mOsm/l) and recovered at room temperature.

Patch-clamp recording were performed using a LNscope (Luigs & Neumann, Germany) equipped with a 40x water immersion objective (Zeiss, Germany), infrared-Dodt gradient contrast and a CMOS camera (Chameleion USB 3.0 monochrome Camera, Point Grey, Canada). Patch pipettes (3-15 MΩ) were pulled from borosilicate glass (GB150F-10, Scientific Products GmbH, Germany) with a horizontal puller (P-1000, Sutter Instruments, Novato, CA, USA). The internal solution contained (in mM): 100 K-gluconate, 20 KCl, 10 Hepes, 4 Mg-ATP, 0.3 Na-GTP, 10 Na_2_-phosphocreatine, 0.3 % biocytine, pH 7.2 (∼300 mosm/l). Data were acquired with an EPC-10 USB amplifier (Heka, Lambrecht, Germany) and Patchmaster Next software (Heka, Lambrecht, Germany). Data were digitized at 20 kHz and lowpass filtered at 10 kHz. Whole-cell patch-clamp recordings consisted of current steps from -100 pA to 300 pA in steps of 50 pA for 500 ms. R_m_ was monitored before and after the cleaning procedure of non-recording pipettes. RMP was monitored during the cleaning process to evaluate any impact of the sonification on the stability of recording. R_pip_, R_GS_ and R_a_ were monitored in voltage-clamp mode to evaluate the efficiency of the cleaning procedure.

### Data analysis

Data were analyzed with Matlab (version 2022a) or Stimfit 0.15 (Christoph Schmidt-Hieber, UCL) and appropriate statistical analysis were performed using GraphPad Prism 8. Representations in Fig. 1B and Fig. 3A were created with BioRender.com.

## Acknowledgments

This work was funded by the Deutsche Forschungsgemeinschaft (DFG, German Research Foundation) Project number 424556709 – GRK 2610 and by Luigs & Neumann GmbH.

## Author contributions

K.J., P.N.R. and B.M.K. conceived the project; K.J. performed the experiments under the supervision of B.M.K.; K.J. and S.W. performed the analysis; K.J. wrote the manuscript with contributions from all authors.

## Data Availability

The datasets generated and analyzed for the current study are available from the corresponding authors on reasonable request.

## References

1. Bezanilla, F., Rojas, E. & Taylor, R. E. Sodium and potassium conductance during a membrane action potential. Journal of Physiology 211, 729–751 (1970).

2. de Hass, V. & Vogel, W. Sodium and potassium currents recorded during an action potential. European Biophysics Journal 1989 17:1 17, 49–51 (1989).

3. Spruston, N. & Johnston, D. Perforated patch-clamp analysis of the passive membrane properties of three classes of hippocampal neurons. J Neurophysiol 67, 508–529 (1992).

4. Petersen, C. C. H. Whole-Cell Recording of Neuronal Membrane Potential during Behavior. Neuron 95, 1266–1281 (2017).

5. Chen, X., Leischner, U., Rochefort, N. L., Nelken, I. & Konnerth, A. Functional mapping of single spines in cortical neurons in vivo. Nature 475, 501–505 (2011).

6. Edwards, F. A., Konnerth, A. & Sakmann, B. Quantal analysis of inhibitory synaptic transmission in the dentate gyrus of rat hippocampal slices: a patchclamp study. Journal of Physiology 430, 213–249 (1990).

7. Bi, G. Q. & Poo, M. M. Synaptic Modifications in Cultured Hippocampal Neurons: Dependence on Spike Timing, Synaptic Strength, and Postsynaptic Cell Type. Journal of Neuroscience 18, 10464–10472 (1998).

8. Kampa, B. M., Letzkus, J. J. & Stuart, G. J. Dendritic mechanisms controlling spike-timing-dependent synaptic plasticity. Trends Neurosci 30, 456–463 (2007).

9. Kampa, B. M., Letzkus, J. J. & Stuart, G. J. Cortical feed-forward networks for binding different streams of sensory information. Nat Neurosci 9, 1472–1473 (2006).

10. Taof, C., Zhangf, G., Xiong, Y. & Zhou, Y. Functional dissection of synaptic circuits: in vivo patch-clamp recording in neuroscience. Front Neural Circuits 9, (2015).

11. Stuart, G. J. & Sakmann, B. Active propagation of somatic action potentials into neocortical pyramidal cell dendrites. Nature 367, 69–72 (1994).

12. Shu, Y., Hasenstaub, A., Duque, A., Yu, Y. & McCormick, D. A. Modulation of intracortical synaptic potentials by presynaptic somatic membrane potential. Nature 441, 761–765 (2006).

13. Seutin, V. & Engel, D. Differences in Na+ conductance density and Na+ channel functional properties between dopamine and GABA neurons of the rat substantia nigra. J Neurophysiol 103, 3099–3114 (2010).

14. Kole, M. H. P. et al. Action potential generation requires a high sodium channel density in the axon initial segment. Nat Neurosci 11, 178–186 (2008).

15. Lee, B. R. et al. Scaled, high fidelity electrophysiological, morphological, and transcriptomic cell characterization. Elife 10, (2021).

16. Cadwell, C. R. et al. Electrophysiological, transcriptomic and morphologic profiling of single neurons using Patch-seq. Nat Biotechnol 34, 199–203 (2016).

17. Li, L. et al. A robot for high yield electrophysiology and morphology of single neurons in vivo. Nat Commun 8, 1–10 (2017).

18. Wang, G. et al. An optogenetics- and imaging-assisted simultaneous multiple patch-clamp recording system for decoding complex neural circuits. Nat Protoc 10, 397–412 (2015).

19. Koos, K. et al. Automatic deep learning-driven label-free image-guided patch clamp system. Nat Commun 12, 1–11 (2021).

20. Neher, E. Ion channels for communication between and within cells. Neuron 8, 605–612 (1992).

21. Kolb, I. et al. Cleaning patch-clamp pipettes for immediate reuse. Sci Rep 6, 1–10 (2016).

22. Kolb, I. et al. PatcherBot: a single-cell electrophysiology robot for adherent cells and brain slices. J Neural Eng 16, 046003 (2019).

23. Davie, J. T. et al. Dendritic patch-clamp recording. Nat Protoc 1, 1235–1247 (2006).

24. Kao, L., Abuladze, N., Shao, X. M., McKeegan, K. & Kurtz, I. A new technique for multiple re-use of planar patch clamp chips. J Neurosci Methods 208, 205–210 (2012).

25. Peng, Y. et al. High-throughput microcircuit analysis of individual human brains through next-generation multineuron patch-clamp. Elife 8, (2019).

26. Ciganok-Hückels, N. et al. Postnatal development of electrophysiological and morphological properties in layer 2/3 and layer 5 pyramidal neurons in the mouse primary visual cortex. Cerebral Cortex (2022) doi:10.1093/CERCOR/BHAC467.

